# Evolution of plasticity in response to ethanol between sister species with different ecological histories (*Drosophila melanogaster* and *D. simulans*)

**DOI:** 10.1101/386334

**Authors:** Sarah A. Signor

**Affiliations:** Molecular and Computational Biology, University of Southern California; Biological Sciences, North Dakota State University

**Keywords:** phenotypic plasticity, *Drosophila*, ethanol, genetic accommodation

## Abstract

The contribution of phenotypic plasticity to adaptation is contentious, with contradictory empirical support for its role in evolution. Here I investigate the possibility that phenotype plasticity has contributed to adaptation to a novel resource. If phenotype plasticity contributes to adaptation, it is thought to evolve in a process termed genetic accommodation. Under this model, the initial response to the environment is widely variable due to cryptic genetic variation, which is then refined by selection to a single adaptive response. I examine the role of phenotypic plasticity in adaptation here by comparing two species of *Drosophila* that differ in their adaptation to ethanol (*Drosophila melanogaster* and *D. simulans)*. Both species are human commensals with a recent cosmopolitan expansion, but only *D. melanogaster* is adapted to ethanol exposure. I measure phenotype plasticity in response to ethanol with gene expression and an approach that combines information about expression and alternative splicing. I find evidence for adaptation to ethanol through genetic accommodation, suggesting that the evolution of phenotype plasticity contributed to the ability of *D. melanogaster* to exploit a novel resource. I also find evidence that alternative splicing may be more important for the adaptive response to ethanol than overall changes in exon expression.

## Introduction

The contribution of phenotypic plasticity to adaptation remains controversial despite considerable empirical and theoretical investigation. This includes whether phenotypic plasticity itself evolves, and how frequently the evolution of plasticity is important for adaptive evolution. For example, is plasticity adaptive, or does it depend upon the cost (if any) of plasticity, and the reliability of cues that induce plastic phenotypes? If plasticity is adaptive, then selection would proceed by favoring the reaction norm that is most adaptive, a process referred to as genetic accommodation (in contrast to simply fixing a different phenotype, known as genetic assimilation) [1,2]. This could potentially accelerate adaptation or divergence because organisms already possess the ability to form alternative phenotypes, rather than requiring the production of a novel phenotype. This is thought to be an important mechanism of adaptation for invasive species, and potentially to be increasingly important in human modified landscapes and climate.

When exposed to novel environments, the predicted path for evolution by genetic accommodation is one in which initially there is increased variance - or genotype-specific differences – in the reaction norm of a trait, followed by refinement by selection for a single adaptive response [1,3–6]. This is because if a trait only exists as an environmentally induced variant, and is therefore not exposed to selection (or infrequently so) then genetic variation should accumulate in the response of that trait to a novel environment [7–10]. After exposure to the novel environment, this cryptic genetic variation is uncovered, manifesting as greater phenotypic variation [11–14]. As lineages adapt to the novel environment, selection on this variation in reaction norm should result in the loss of variation for this trait as genetic accommodation occurs [15]. As long as the environment is heterogeneous the expectation is that it will remain a plastic trait, rather than evolving to be stably expressed. This would manifest as a transient increase in genotype by environment interactions, followed by reduced genotype by environment interactions but persistent environmental response, a pattern which is consistent evidence of adaptation [16–20].

While changes in heritability or variance have been previously shown between populations occupying ancestral and novel habitats, inbred lines have not been used previously to specifically measure genotype by environment interactions in adapted and non-adapted populations. Furthermore, in general the evolution of phenotypic plasticity is investigated with regards to an ancestral and a novel environment, rather than a heterogeneous environment where plasticity is expected to be maintained. In this manuscript I will test the predictions from genetic accommodation using inbred lines in *Drosophila*, an approach which will allow direct quantification of the genotype by environment interaction. I will also investigate a heterogeneous environmental variable, ethanol, for which different species of *Drosophila* have different adaptive histories.

*D. simulans* and *D. melanogaster* have divergent adaptive histories with respect to ethanol. *D. melanogaster* is considerably more ethanol tolerant than *D. simulans*, and is found regularly feeding upon and ovipositing in resources with ethanol concentrations greater than 8% [21–23]. At concentrations of 4% ethanol *D. simulans* shows reductions in survivorship and increased development time relative to *D. melanogaster* [24]. Furthermore*, D. melanogaster* and *D. simulans* are both cosmopolitan species and human commensals, however while *D. melanogaster* is commonly found inside houses, breweries, and wineries, *D. simulans* is more often observed in orchards and parks (though their niches overlap significantly and they are often found on the same patches) [24–33]. Therefore, the expectation is that *D. simulans* will show a non-adapted phenotype manifested as considerable genotype by environment interaction. *D. melanogaster* will retain an interaction with the environment, but it will not be genotype-specific, which is consistent with adaptation to a heterogeneous environment.

The trait I measure to test this prediction is gene expression. Phenotypic plasticity is often achieved by dynamic changes in gene expression across environments, and current evidence suggests that it plays an important role in regulating evolutionarily important phenotypes [34–37]. Gene by environment interactions also appear to be a common attribute of gene expression, and yet while many studies have quantified the effect of the environment on gene expression, far fewer have performed these experiments in multiple genotypes to obtain the interaction between genotype and the environment for gene expression [38]. Indeed, most work on gene expression plasticity has not quantified the interaction between genotype and the environment [36,38–42].

Short read sequencing fundamentally cannot separate the contributions of alternative splicing and gene expression to changes in abundance of an exon, thus I approach the analysis in a way that includes both phenomenon (gene expression and alternative splicing, see Methods). In addition, because of the potential for alternative splicing to respond more rapidly to environmental differences than gene expression *sensu stricto*, it may be one of the earliest mediators of environmental response. This is because it does not require de novo transcription or protein production. It is thought to diverge more rapidly than gene expression between lineages, and it has been linked to stress responses and changes in transcriptome complexity in response to environmental differences [43–50]. Furthermore, it has been previously implicated in the response to ethanol, and in *D. melanogaster* may be a more important component of the response than gene expression changes [51–57]. Yet it remains the topic of few investigations into environmental response.

Here I investigate plasticity in response to ethanol, including genetic variation for plasticity over time, in *D. simulans* which is not adapted to ethanol use. I will compare these results to previous work on *D. melanogaster*. I find that in *D. simulans* there is an order of magnitude larger effect of genotype by environment compared to *D. melanogaster*, however both species maintain an environmental response of similar magnitude. This suggests the following scenario: In *D. simulans* high ethanol concentrations are essentially a novel environment that reveals cryptic variation for phenotypic plasticity. In *D. melanogaster*, genetic variation in phenotypic plasticity has been removed by selection in favor of a single environmental response through a process of genetic accommodation. This single environmental response involves more alternative splicing than observed in *D. simulans*, suggesting it is an important component of adaptation.

## Methods

### Fly lines

Six genotypes were used in these experiments. The genotypes used for male flies from *D. simulans* were collected from the Zuma Organic Orchard in Zuma Beach, CA in the winter of 2012 and made nearly isogenic by 15 generations of full sib inbreeding [58,59]. In natural conditions flies will not by homozygous at all loci, thus each inbred genotype was crossed to a white eyed ‘tester’ strain to create the F1 flies used in the ethanol exposure assays (*D. simulans*, *w^501^*, Cornell species stock number 14021-0251.011). Rearing occurred on a standard medium at 25 °C with a 12-h light/12-h dark cycle. In order to control for maternal effects and variation in offspring quality, female parents were collected as virgins, aged one day, and then density matched with male flies (10 per sex). The F1 offspring were then collected as virgins, reared in single sex vials with standardized density (24-30 flies if male, 12-15 if female), and aged for three to four days prior to the assay. A portion of this experiment was intended to analyze behavioral differences in ethanol, thus during the ethanol exposure a female was included as a stimulus, but was not collected for RNA-seq [59,60]. The day prior to the ethanol exposure these females were mated to flies from a standard genotype, such that in the final assay each chamber contained one mated female fly from the *y^1^w^1^* mutant lines and two virgin males from the target F1 cross, of which only the latter were collected for RNA-seq. All procedures used here in the maintenance and raising of *D. simulans* are the same as those used previously in *D. melanogaster*, facilitating easy comparison between studies [59,60].

### Experimental setup

In this experiment intoxication occurs through inhalation of ethanol vapors, and evidence of the behavioral effects in both species and the efficacy of this approach have been published previously [59,60]. Ethanol exposure took place in a circular arena, each of which was part of a larger chamber containing 12 arenas each with a diameter of 2.54 cm (VWR cat. no. 89093-496) (Figure 1). The bottom of each chamber contained a standard amount of either grapefruit medium or medium in which 15% of the water has been replaced with ethanol. Replicates for both species were conducted randomly under standardized conditions (25 °C, 70% humidity). Prior to the assay the flies were sedated through exposure at 4 °C for ten minutes, to avoid the confounding effects of CO_2_, then placed in the chambers which were left then upside down at room temperature for ten minutes while the flies oriented themselves [59–61]. After ten minutes the timing of the assay began, and were conducted for ten, 20, or 30 minutes for three replicates of each of the two conditions (Figure 1). Flies are most active during the hours following dawn, thus to standardize behavior and circadian rhythms all assays were conducted within a two-hour window after dawn. At the conclusion of the assay the chambers were flash frozen in liquid nitrogen, allowed to freeze through, transferred to dry ice, and all of the males were collected for RNA-seq.

**Figure 1:**
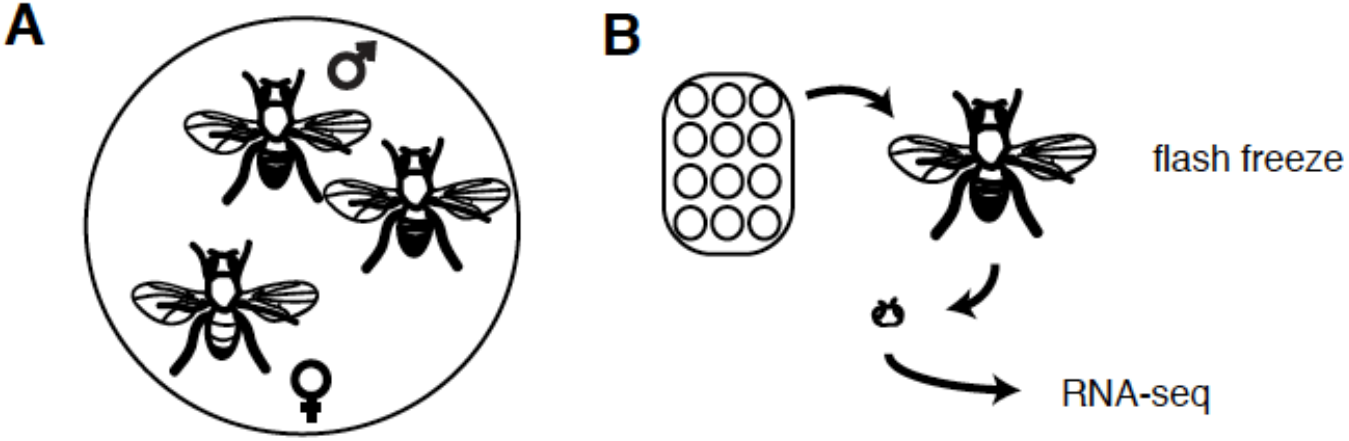
Experimental setup. A. This is an illustration of the social environment that each *Drosophila* male was exposed to during the experiment. Each chamber contained two male flies and one female fly. Within a plate there were twelve chambers, and the males from a single plate where pooled to create a sequencing library. B. Males from each of the twelve chambers were collected and flash frozen after either 10 minutes, 20 minutes, or 30 minutes. After flash freezing their heads were isolated for RNA-seq.

**Figure 1:**
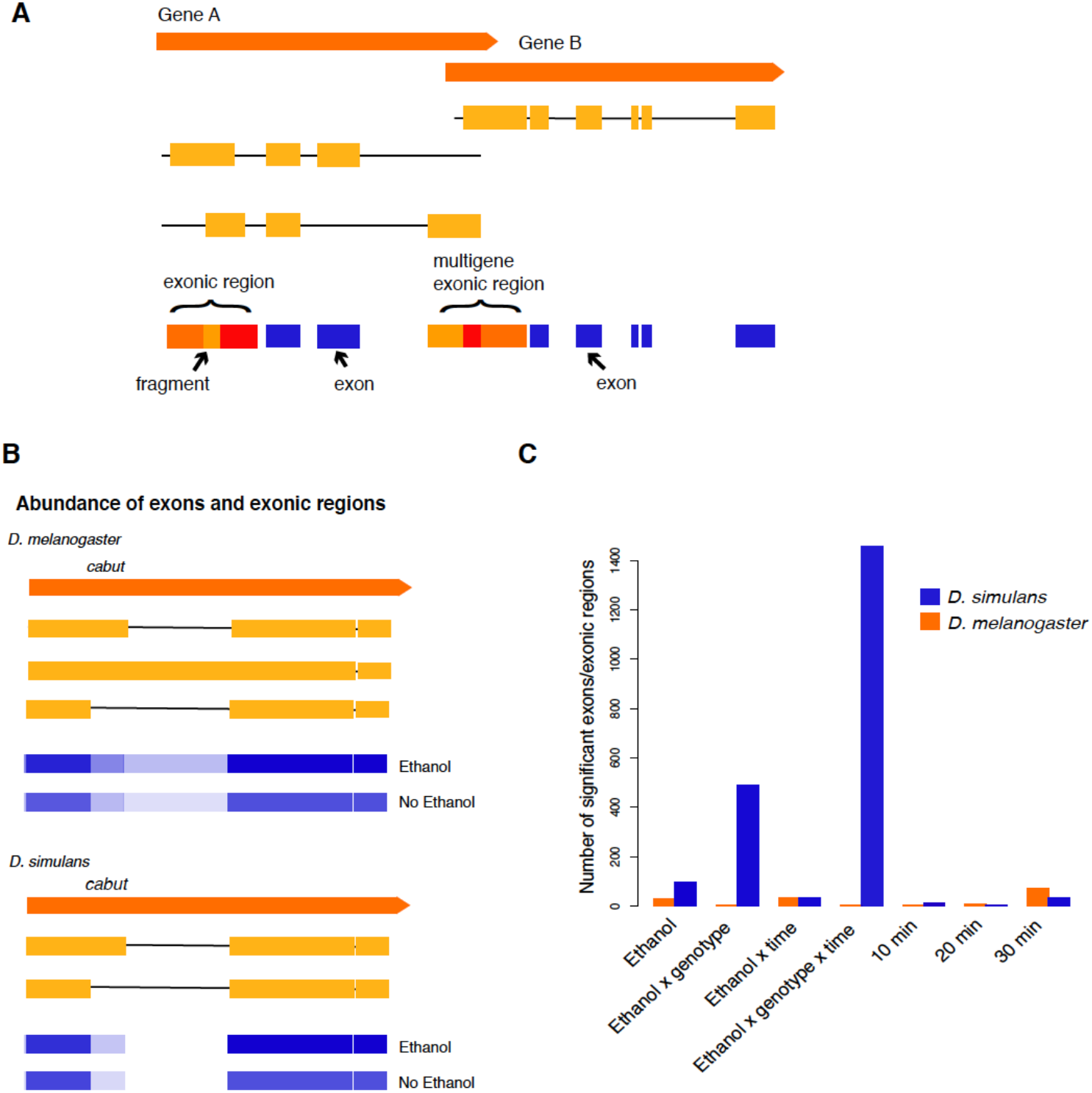
**A**. An illustration of the classification scheme for exons, exonic regions, and exon fragments. Exons either do not overlap other exons in different isoforms (exons, shown in blue), or are fused regions consisting of a set of overlapping exons (exonic regions, shown in shades of orange and red). Exonic regions can be decomposed into exon fragments, depending upon their overlap between different isoforms. **B.** An example of the information about gene expression and isoform abundance that can be inferred from measures of differential exon abundance. *cabut* is a relatively simple example, having few isoforms and exons, and yet it is still a complicated inter-species comparison given that *D. melanogaster* has more annotated isoforms that *D. simulans*. Below each gene model the frequency of different exons and exon fragments in each environment is shown, with the most frequent in dark blue. In *D. simulans* the first of two isoforms shown has an exon fragment that is more abundant in the presence of ethanol, suggesting that that isoform is more abundant. In *D. melanogaster* only the second of the three isoforms has a unique fragment, which is also more abundant in ethanol and suggests differential isoform usage. **C.** The number of exons or exonic regions that are significant for each component of variance in *D. melanogaster* and *D. simulans*. Many more exons and exonic regions are significant for *D. simulans* than *D. melanogaster*, but only for components of variance that contain an interaction with genotype

### Sample preparation and RNA sequencing

Flash-frozen flies were freeze dried and ten to 12 heads were placed into a 96-tube plate (Axygen MTS-11-C-R). mRNA purification, cDNA synthesis and library preparation were carried out by RAPiD GENOMICS (http://rapid-genomics.com) using a robot. mRNA was purified using Dynabeads mRNA DIRECT Micro kit (Invitrogen # 61021) with slight modifications. To fragment the RNA mRNA-beads were resuspended in 10 uL 2X first strand buffer (Invitrogen # 18064-014), incubated at 80 C for two minutes and placed on ice, then the supernatant was collected after five minutes on a magnetic stand. First strand synthesis was performed using standard protocols for Superscript II (Invitrogen #18640-014) and reverse transcription (25 C 10 min, 42 C for 50 min, 70 C for 15 min, 10 C hold). Second strand synthesis was carried out using standard protocols with DNA Pol I and incubated at 16 C for 2.5 hours. cDNA was purified with 1.8 volume of AMPure XP following manufactures instructions (Beckman Coulter A63880). Sequencing was performed using the Illumina HiSeq 2500 as both 2 × 150 bp or 2 × 50 bp reads, resulting in two technical replicates per sample. The two run lengths (and runs) were intended to provide extra coverage, and all replicates were sequenced in both runs.

### Exon expression analysis

It is common in organisms with alternative splicing for exons from different isoforms of a single gene to overlap with one another, or be shared between all or most isoforms (Figure 1A). Short read data fundamentally cannot resolve these exons to individual isoforms, however, one approach is to quantify each exon separately and decompose exons overlapping between isoforms into those which are shared and unique. When the differences between overlapping exons are less than 10 bp, there is no appreciable amount of information loss in not decomposing overlapping exons, and this approach has been taken in the past [62–65]. However, in many cases the differences in exon overlap are much larger than this, so to address this issue I use a classification scheme where reads may be assigned to *exons*, *exonic regions*, or *exon fragments*, and then compare the abundance of each exon/exonic region/exon fragment in each condition (i.e. the abundance of an exon with and without ethanol) [61] (Figure 1A). Here, an *exon* does not overlap any other exons. If exons from different isoforms overlap, they are grouped into an *exonic region*. *Exon fragments* are classified by decomposing *exonic regions* based on the 5’ and 3’ positions of the exons within the region. Thus, all *exon fragments* are subregions of *exonic regions*. Exon boundaries were determined using the *D. melanogaster* FlyBase 6.17 genome features file and the *D. simulans* 2.02 genome features file. Alignment was performed using BWA-MEM version 0.7.15 (which has been shown to perform better than split read mappers such as STAR [66,67]) and BED files were used to count reads in each region and obtain the length adjusted read count (reads in region divided by the length of region), and the APN (average per nucleotide) [68].

The APN was calculated separately for each read length and then combined between read lengths to handle the mixture of read lengths for each sample (2 × 150 bp and 2 × 50 bp). If the APN was greater than zero in at least half of all samples per condition it was considered detected. While I considered several approaches to normalize coverage counts upper-quartile normalization with log-transformation and median centering within time × treatment × genotype were selected due to better performance of the residuals [69,70].

To test the significance of components of expression variation, the log APN for each exonic region was modeled as

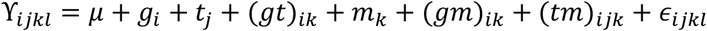

for the *i*^th^ genotype (*g*_*i*_), *j*^th^ treatment (*t*_*j*_; *j* = ethanol or no ethanol), *k*^th^ time point (*m*_*k*_; *k* = 10 min, 20 min, 30 min), and *l*^th^ replicate (Supplemental File 1-2). For the interaction between treatment and time point, the log APN for each exonic region was modeled as

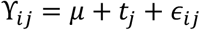

or the *i*^th^ condition (time × treatment) and *j*^th^ replicate. Contrasts to compare treatments within time point (ethanol versus no ethanol, for ten, 20 and 30 minutes) were conducted (Supplemental Files 1-2). Residuals were evaluated for conformation with normality assumptions, and assumptions were met in excess of 95% of the models.

In contrast to exon abundance, I evaluated alternative splicing specifically by comparing the abundance of all exons within a gene to each other, as this is more direct evidence of a change in isoform abundance than a change in exon/exonic region/exon fragment abundance; Figure 2A). For example, a change in relative abundance might be detected if the last exon of a gene was the most abundant compared to all other exons/exonic regions/exon fragments without ethanol and the least abundant with ethanol. Exons and exonic regions for each gene and for each sample were ranked and the most expressed region ranked as one, the least expressed region as three and all others as two (Figure 2A). Exon ranks for each gene were modeled as

**Figure 2:**
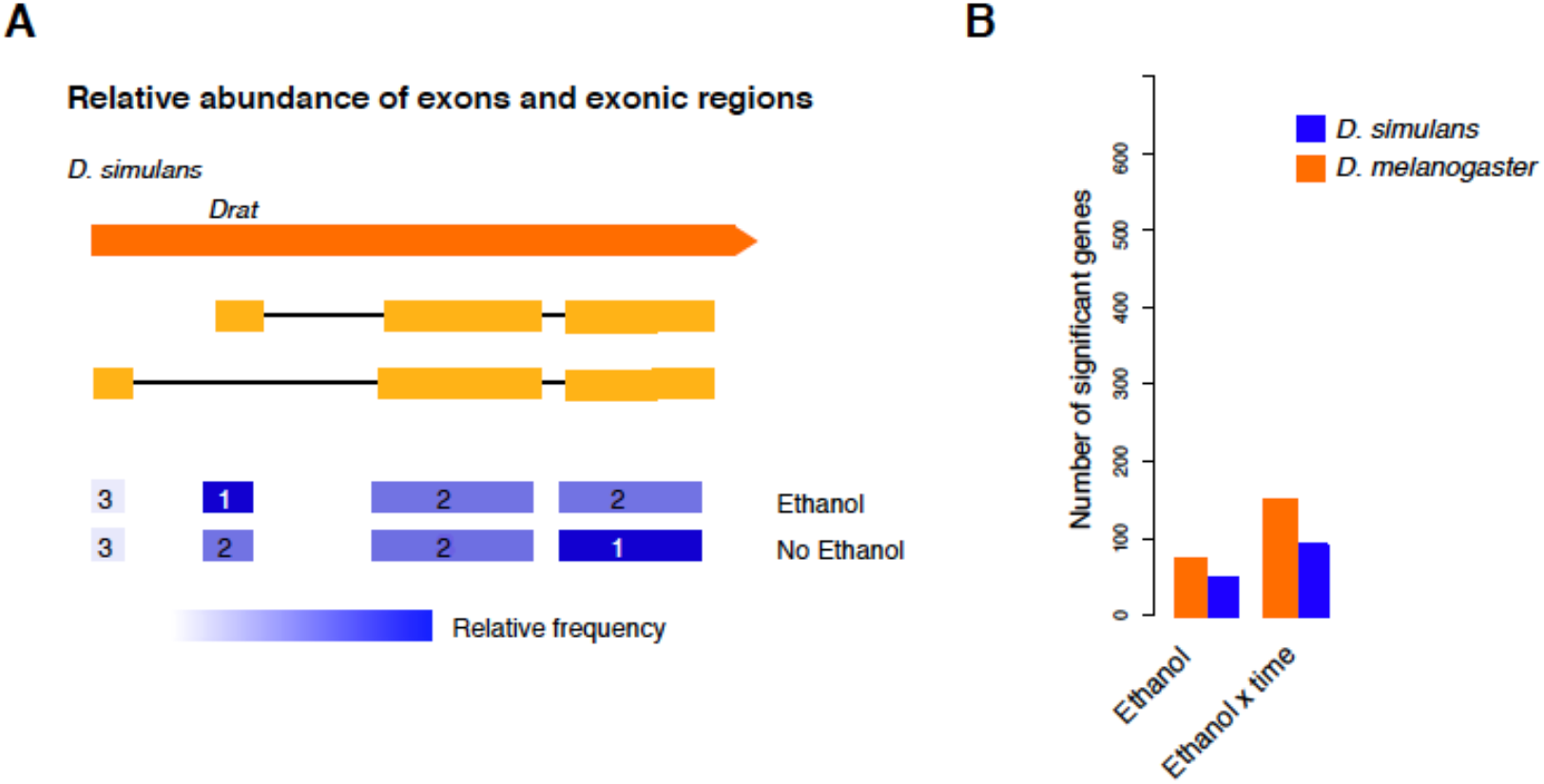
**A.** An illustration of a change in the relative abundance of the gene *Drat*. Only the example of *D. simulans* is shown. With and without ethanol different exons are the most abundant compared to the other exons within the gene. As one of these exons is not shared among all isoforms, this suggests a change in isoform usage between environments. **B.** The number of genes that have a significant change in relative abundance for each component of variance in *D. melanogaster* and *D. simulans*.

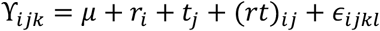

where Υ_*ijk*_ is the exon rank (1,2,3) of the *i*^th^ exonic region of the gene, *j*^th^ condition (time × treatment), and the *k*^th^ replicate; *r*_*i*_ is the exonic region of the gene; *t*_*j*_ is condition; and (*rt*)_*ij*_ is the interaction between exonic region and condition. More traditional GLM approaches can only be taken if their assumptions are met, and in this case they are not due to a lack of normality in the distribution of model residuals. Accordingly, a non-parametric test must be relied upon to look for changes in exon or exonic region representation between exons of a gene and I used a rank test to summarize changes in exon representation (Supplemental File 3). F-tests for the significance of the mean square attributed to the effect tested versus the mean square attributed to error, or the appropriate interaction term, were used. The false discovery rate was controlled using the Benjamini-Hochberg procedure, with a significance cutoff of α = 0.05 [71]

### GO Analysis

When a gene in *D. simulans* had more than one ortholog in *D. melanogaster* only one ortholog was included for the GO analysis, so as not to inflate the number of genes involved in a given process. This does presume that orthologs will be annotated with the same GO terms, and this is generally the case. For example, in *D. simulans* there is only *AOX4*, while in *D. melanogaster* there is *AOX3* and *AOX4*, but the GO terms for each are the same. However, as *D. simulans* genes are generally not independently annotated, especially those without *D. melanogaster* orthologs, if there was no *D. melanogaster* ortholog the gene was not included in the enrichment analysis. Lists of significant genes were tested for GO enrichment using the PANTHER classification system [72]. They were corrected for multiple testing and a *p*-value of 0.01 was required for significance.

### Functional class enrichment

I tested the significant sets of exons and exonic regions for enrichment with non-protein coding genes. Multigene exonic regions were not included, meaning exons that belong to more than one gene, as they often do not correspond to the same functional class of gene. Every test of functional class enrichment compared the frequency of a given subcategory among all exons and exonic regions detected in the dataset compared to the frequency within a significant list of exons and exonic regions. A χ^2^ test was performed in R to test the significance of the enrichment of each of these categories.

## Results

### Exon expression and isoform usage

It is difficult to decouple alternative isoform usage from gene expression, given that many exons are shared between isoforms or overlap other exons. To infer isoforms from short read data, one must rely upon unique junctions or regions of individual isoforms and extrapolate to shared regions. This requires accurate isoform annotation (knowing that any given exon/junction is found in combination with other exons/junctions) and in general can be very noisy. Accordingly, I subscribe to a simpler but more robust approach and summarize the abundance of different exons and exonic regions separately [61,62,64,65]. I detail overall abundance of exons, exonic regions, and exon fragments, and changes in the relative abundance of exons within a gene (Figure 2 A-C; Figure 3 A&B).

### Changes in exon expression in D. simulans

Table 1 summarizes the number of exons, exonic regions, and exon fragments which alter their expression in response to ethanol for *D. simulans*, and a full list of genes is available in the Supplemental File 1 & 2. Compared to previous work in *D. melanogaster*, for treatment seven exons and exonic regions were shared between species: *Drat*, *cabut*, *CG11741*, *CG32512*, *CG4607*, *Pinocchio*, and *sugarbabe* (Table 1; Figure 2 A-C, [61]). For the interaction between ethanol and time three genes were shared between exons and exonic regions for these species, *Drat*, *cabut*, and *CG43366*. At ten and twenty minutes *cabut* is shared between exons and exonic regions in *D. melanogaster* and *D. simulans*. At 30 minutes *Drat*, *CG32103*, *sugarbabe*, *cabut*, *Pinocchio*, *CG32512*, and *CG4607* are shared between species. This suggests that *Drat*, *cabut*, *sugarbabe*, *Pinocchio*, *CG32512*, and *CG407* are important for the response to ethanol, as they are shared between species for multiple components of variance and have been implicated in the response to ethanol previously [73–75].

**Table 1:** A summary of the exons, exonic regions, and exon fragments that are significantly differently expressed for each component of variance. Abbreviations are as follows: (G) Genotype, (ETOH) Ethanol, and (T) Time. Ethanol in this case is the environment.

|  | G | ETOH | G x ETOH | ETOH x T | G x ETOH x T | 10 | 20 | 30 |
| --- | --- | --- | --- | --- | --- | --- | --- | --- |
| Exons | 1994 | 76 | 387 | 24 | 1158 | 9 | 3 | 29 |
| Exonic Regions | 406 | 23 | 98 | 7 | 299 | 3 | 1 | 4 |
| Fragments | 608 | 18 | 81 | 6 | 225 | 4 | 2 | 6 |

*D. simulans*

*D. simulans* is enriched for several GO terms in response to interactions with genotype, including several terms relating to cilia, such as the axonemal dynein complex and ciliary part (Table 2). GO terms were also enriched for the sensory perception of taste and detection of chemical stimulus. Many components of variance in *D. simulans* are not enriched for GO terms due in part to the frequency with which non-protein coding genes were implicated – for example in response to ethanol 15% of the implicated genes do not have a *D. melanogaster* ortholog, and of those only two are protein coding.

**Table 2:** A summary of the significant GO enrichment terms for *D. simulans*. The abbreviations are the same as in Table 1. *p*-values are listed adjacent to the significant term, the significance cutoff is *p* < .01.

|  | Biological Process |  | Cellular Component |  | Molecular Function |  |
| --- | --- | --- | --- | --- | --- | --- |
| *G x ETOH* | | | axonemal dynein complex | $3 \times 10^{-3}$ | dynein light chain binding | $3.3 \times 10^{-3}$ |
| *G x ETOH x T* | sensory perception of taste | $6.6 \times 10^{-3}$ | nucleoplasm | $6.5 \times 10^{-3}$ | RNA binding | $7.8 \times 10^{-3}$ |
| | detection of chemical stimulus | $8.2 \times 10^{-3}$ | cytoskeletal part | $8.4 \times 10^{-3}$ | | |
| | | | ciliary part | $4.6 \times 10^{-3}$ | | |

### Changes in the relative abundance of exons within a gene in D. simulans

In response to treatment in *D. simulans* 54 genes showed changes in the relative abundance of their exons/exonic regions, and 94 genes change the relative abundance of their exons for the interaction between treatment and time (Figure 3B, for a full list see Supplemental File 3). For the interaction between treatment and time *lola*, *Mhc*, and *Prm* were shared between species. Changes in relative abundance were not enriched for any GO terms in *D. simulans*, with the exception of peroxiredoxin activity in response to ethanol, albeit slightly above the more conservative cut-off used in my other tests (*p* = .017). Peroxiredoxin activity has been associated with protection against alcohol induced liver damage [76,77].

#### In D. simulans genotype-specific reactions to the environment are abundant

In *D. simulans* many exons and exonic regions are significant for components of variance that interact with genotype: 1457 for the interaction between genotype, treatment and time, and 486 for the interaction between genotype and treatment, compared to two and three exons and exonic regions respectively in *D. melanogaster* (Table 1; Figure 2 A-C). For example, for the interaction between genotype, treatment and time. 1% as many exons were implicated in *D. melanogaster* as in *D. simulans*. For other components of variance, including genotype (suggesting that this is not due to differences in overall polymorphism), the number of exons and exonic regions implicated in expression differences is comparable between species. For example, for genotype 87% as many exons were implicated in *D. melanogaster* as in *D. simulans*; for the interaction between ethanol and time it is 97%; to highlight the scale of the difference.

#### In D. simulans interactions with genotype are enriched for nested non-protein coding genes

A large number of the genes implicated in differences due to interaction with genotype were non-protein coding genes. The number of non-protein coding genes that were significant for ethanol, ethanol by genotype, and ethanol by genotype by time, were more than would be expected by chance in *D. simulans* (χ^2^ = 49.598, *p* < 0.0005; χ^2^ = 235.21, *p* < 2.2 × 10^−16^; χ^2^ = 727.17, *p* < 2.2 × 10^−16^). These are generally long non-coding RNAs (lncRNAs) as the shortest is annotated at 713 bp, and the majority are over 4,000 bp (long non-coding RNAs being any non-protein coding genes over 200 nt), with some pseudogenes. *D. melanogaster* is not enriched for non-protein coding genes in response to ethanol, ethanol by time, ethanol by genotype, ethanol by genotype by time, ten or 20 minutes, exons or exonic regions expressed only in one environment, or genes implicated in changes in relative abundance [61]. However, at 30 minutes *D. melanogaster* is enriched for non-protein coding genes, though this concerns far fewer genes than in *D. simulans* (χ^2^ = 12.831, *p* = 0.005).

Many of the non-protein coding genes implicated in interactions with genotype are nested in the introns of other genes, thus I sought to determine if it was more than expected by chance. First, I must determine the overall frequency of gene nesting, using the criteria that an exon nested in an intron must overlap the intron by at least 80 bp or 10%. I found that in *D. melanogaster* 9.2% of exons were nested within introns, while in *D. simulans* 9.7% were nested, similar to what has been previously reported [78]. This was reflected in the data, where for both *D. melanogaster* and *D. simulans* 6.8% of exons and exonic regions were nested within introns (lower because multi-gene exons were excluded). However, among significant exons and exonic regions in *D. melanogaster* 18.6% were nested, while in *D. simulans* 33.5% were nested [61]. This is a significant enrichment of exons that are nested within other introns, for both *D. melanogaster* (χ^2^ = 12.344, *p* < .0004) and *D. simulans* (χ^2^ = 1721.1, *p* < 2.2 × 10^−16^). In addition, compared to the total number of nested genes that are noncoding within the dataset, the number that are significant for the response to ethanol is enriched for nested, non-protein coding genes in *D. simulans*(χ^2^ = 237.49, *p* < 2.2 × 10^−16^), but not in *D. melanogaster*. Of these nested non-protein coding genes, 83% are on the opposite strand as their parental genes, suggesting that these genes are regulated independently of their parental genes (and their expression is uncorrelated, see Supplemental File 4). Overall *D. melanogaster* has more annotated non-protein coding genes (2963) than *D. simulans* (1675), and similarly more exons from noncoding genes are nested within introns (1772 *D. melanogaster*, 1066 *D. simulans*). The importance of nested non-protein coding genes for the response to the environment, or potentially delays in splicing that are specific to certain components of variance, is unclear.

## Discussion

The prediction from theory on genetic accommodation was that in *D. simulans* there would be abundant cryptic genetic variation in response to ethanol exposure, while in *D. melanogaster* there would be a response to the environment that was not genotype-specific. This is because *D. simulans* does not exploit ethanol-rich resources, thus environmentally induced variants are not exposed to selection and accumulate as cryptic genetic variation. This manifests as greater variation between genotypes in the novel environment [11–14]. In *D. melanogaster* ethanol is used as a resource, however the environment is patchy and therefore response to ethanol is expected to be selected upon for the optimal response, but to remain plastic. This would manifest as an environmental response, but without extensive genotype-specific variation for that response [16–20]. The use of inbred lines is a unique opportunity to assess these predictions, as the contribution of interactions with genotype can be directly assessed.

Here I show that these expectations are met in *D. melanogaster* and *D. simulans*. In *D. melanogaster* adaptation to ethanol has occurred through genetic accommodation – selection on plasticity to reduce genetic variation and produce a single plastic response. This has facilitated expansion to a novel resource in *D. melanogaster*, ethanol rich substrates. In *D. simulans* this is not the case, and lack of selection in ethanol environments has allowed for the accumulation of cryptic genetic variation for phenotype plasticity manifested as extensive genotype by environment interactions. Joshi and Thompson (1997) found that in *D. simulans* there was a reduction in variation between families for the phenotypic response to ethanol substrate after selection for tolerance to ethanol. This is consistent with the observed scenario in D*. melanogaster* and *D. simulans* – adaptation to a novel environment results in a reduction in variation for environmental response and while retaining a single response to the environment. This is also consistent with genetic accommodation as a mechanism of adaptation to ethanol, where initial responses to environmental differences are a mix of adaptive and maladaptive, which are subsequently honed by selection.

Previous literature on the response to novel environments is equivocal concerning the expectation that plasticity evolves from a starting point of increased genetic variance due to cryptic genetic variation.

For example, early estimates of heritability found reduced expression of genetic variation under ‘stressful’ conditions – though stressful does not necessarily imply novel [79,80]. Work in *Drosophila* has been similarly ambiguous – in *D. mojavensis* a similar reduction in genotype by environment interactions was detected in adaptation to a novel cactus host, however in *D. melanogaster* differences in genotype by environment interactions were not detected between cold-adapted and non-cold adapted populations [16,42]. These differences may be due to difficulty in defining novelty, and in equating novelty with stress. As posited by Chevin and Hoffman (2017), it may be that genetic variance increased in novel environments, but that favorable conditions are rare – meaning that ‘stressful’ environments are commonly experienced resulting in a reduction in genetic variation.

The abundance of lncRNAs which are involved in genotype by environment interactions in *D. simulans* is perhaps not surprising, given that they have been implicated previously in dynamic responses [81]. It is also possible they are more frequently involved because lncRNAs are often less conserved, as there is no requirement for the maintenance of ORFs and codon synonymy [82–85]. If neutral variation was allowed to accumulate without selection, and then uncovered in a novel environment (cryptic genetic variation), preferential accumulation within less constrained sequences would be expected.

Alternative splicing has the potential to be an important component of adaptation and response to the environment, as it does not require de novo transcription or protein production. In *D. melanogaster* it is a more important component of the response to ethanol than in *D. simulans*, as evidenced by differences in the frequency of significant changes in relative abundance of exons (nearly twice as many in *D. melanogaster* [61]). The response to ethanol in *D. melanogaster* is adaptive, thus it can be deduced that alternative splicing is important for adaptation to ethanol. Alternatively, the fact that it is less important in *D. simulans* suggests that perhaps there is selection against the accumulation of cryptic genetic variation for splicing patterns. This is not surprising – previous studies have linked alternative splicing to the response to the environment [43–50], as well as important adaptive differences [43,86–88].

Furthermore, it has been previously implicated in the response to ethanol, and in *D. melanogaster* may be a more important component of the response than gene expression changes [51–57].

In general, the GO terms implicated in this study relate to known effects of ethanol, for example, alcohol-induced ciliary dysfunction is a known consequence of alcohol exposure [89,90]. Furthermore, the genes implicated in both species (*Drat*, *cabut*, *sugarbabe*, *Pinocchio*, *CG32512*, and *CG407*) have generally been implicated in the response to ethanol in other studies. *cabut* has been previously observed as being upregulated in response to ethanol, and it is in general responsive to changes in metabolic conditions [75]; [91,92]. *sugarbabe* and *cabut* are both downstream of the *Mondo-Mlx* sugar sensing pathway [93], which has been linked to severe obesity, high circulating triglycerides, and tumorigenesis [94–97]. *Drat* has been implicated in ethanol-related cell death [98]. Specifically with regard to other studies of ethanol exposure in *Drosophila* overlap is low between significant datasets, however this may in large part be due to differences in methodology [61].

Inferring that gene expression differences are adaptive or non-adaptive remains a major challenge in the study of gene expression reaction norms, given the lack of direct correlation between gene expression phenotypes and organismal phenotypes. However, the patterns observed in *D. simulans* suggest that abundant genotype by environment interactions have accumulated neutrally and become uncovered in response to a novel environment. In contrast, in *D. melanogaster* this ethanol environment is not novel and variation in plasticity has been selected out in favor of an adaptive phenotypic response. lncRNAs are preferentially differentially expressed in *D. simulans* in response to ethanol likely because they are less constrained and can accumulate more neutral variation. This is an excellent illustration of genetic accommodation, where phenotypic plasticity has encouraged adaptation to a novel resource within a patchy environment.

## Acknowledgements

I would like to thank Sergey Nuzhdin, Jeremy Newman, and Lauren McIntyre for contributions to the manuscript. I would like to thank my undergraduate researchers for assistance in producing this data: N. Shadman, V. Paterson, Z. Polonus, K. Cortez, L. Hassanzadeh, L. Cline, A. Khokhar, E. Lee, K.L. Yee, M. Ling, S. Sarva, O. Akintonwa, A. Gupta, R. Manson, P. Hassanzadeh, and K. Kavoussi.

## Data Accessibility

*D. melanogaster* sequence data have been submitted to Bioproject: accession number PRJNA482662. *D. simulans* sequence data will be made available upon acceptance.

